# Harmonized diffusion MRI data and white matter measures from the Adolescent Brain Cognitive Development Study

**DOI:** 10.1101/2023.04.04.535587

**Authors:** Suheyla Cetin-Karayumak, Fan Zhang, Tashrif Billah, Leo Zekelman, Nikos Makris, Steve Pieper, Lauren J. O’Donnell, Yogesh Rathi

## Abstract

The Adolescent Brain Cognitive Development (ABCD) study has collected data from over 10,000 children across 21 sites, providing valuable insights into adolescent brain development. However, site-specific scanner variability has made it challenging to use diffusion MRI (dMRI) data from this study. To address this, a database of harmonized and processed ABCD dMRI data has been created, comprising quality-controlled imaging data from 9345 subjects. This resource required significant computational effort, taking ∼50,000 CPU hours to harmonize the data, perform white matter parcellation, and run whole brain tractography. The database includes harmonized dMRI data, 800 white matter clusters, 73 anatomically labeled white matter tracts both in full-resolution (for analysis) and low-resolution (for visualization), and 804 different dMRI-derived measures per subject. It is available via the NIMH Data Archive and offers tremendous potential for scientific discoveries in structural connectivity studies of neurodevelopment in children and adolescents. Additionally, several post-harmonization experiments were conducted to demonstrate the success of the harmonization process on the ABCD dataset.

## 1) Background and Summary

The Adolescent Brain Cognitive Development (ABCD) Study® is a landmark research project focused on child health and brain development ^1^. The ABCD Research Consortium consists of 21 research sites across the US and has collected one of the largest multi-domain datasets, including neuroimaging, behavioral, cognitive, and genetic data, from over 10,000 children from middle childhood into early adulthood. Using this extensive dataset, the ABCD study aims to provide the opportunity to understand the factors that shape brain and cognitive development during this crucial period of life. To ensure uniformity in scanning protocols, the ABCD study has taken extensive measures to maintain imaging acquisition parameters and protocols consistent across all 21 participating sites ^2^. Despite these efforts, data acquired from different sites and vendors (Siemens, Philips, and GE) introduce significant site effects ^3^, due to scanner-specific MR acquisition sequences and data reconstruction algorithms. In fact, given the vast array of design choices and vendor-specific decisions beyond the users’ control, it is almost always challenging to completely standardize the different scanners in multi-site neuroimaging studies such as the ABCD study, leading to inter-scanner differences in the acquired data.

Diffusion-weighted magnetic resonance imaging (dMRI) is a non-invasive imaging technique that can detect microstructural changes and reveal white matter connectivity in the brain. Studies have shown that even small scanner-related differences in the dMRI data collected from various scanners can result in significant measurement biases in the dMRI measures of white matter connectivity and microstructure ^4–10^. Even when scanners from the same manufacturer are used across different sites, the measurement biases in the dMRI data can be substantial. For example, Thieleking et al. ^11^ showed that the difference in the fractional anisotropy (FA) from the same healthy participants (N = 121) scanned on two 3T Siemens Magnetom scanners could be up to 33 times larger than the effect seen in healthy aging. In addition, Schilling et al. ^12^ demonstrated the significant variability in connectivity measures across different scanners (e.g., the density of streamlines in white matter fiber connections/tracts), which also resulted in significant differences in the related microstructural measures. Moreover, multiple studies have highlighted that scanner-related variability is highly non-linear across various tissues and white matter regions/tracts ^6, 9, 10, 12–14^. Thus, many studies underscore the significance of eliminating bias due to scanner or site from dMRI data at the voxel level before running joint analysis in multi-site studies such as the ABCD study ^3, 5, 6, 15–20^.

Enabling pooled large-scale analysis of multi-site datasets represents an outstanding opportunity for improving our understanding of the human brain in health and disease. However, such research requires large sample sizes, which can only be obtained by appropriate pooling and removal of scanner-specific effects from dMRI data acquired from multiple scanners. *“Harmonization”* is a way to mitigate the measurement differences attributed to the scanner-, protocol-, or other site-related effects ^5, 6, 15–20^. The goal of harmonizing dMRI data is to preserve variability that is purely related to biology or disease and remove variability caused by intrinsic or acquisition-related factors of the scanners, which can conceal the desired effect. Various data harmonization methods have been proposed in the literature ^7, 18, 21–25^. A majority of these approaches are based on adding statistical covariates to remove site effects. These methods are commonly used in multiple fields, from genetics to functional MRI and recently in dMRI ^18, 21–25^. These methods typically involve fitting a dMRI model (such as diffusion tensor imaging) to obtain the desired dMRI measures (e.g., FA) for each subject at each site, followed by regression modeling where the site is added as a linear covariate to minimize inter-site effects. These approaches are limited for several reasons. First, they assume linear site effects in the microstructural measures (such as FA, mean diffusivity, kurtosis, etc.). However, this assumption has been challenged by many studies demonstrating nonlinear site effects ^6, 14, 26, 27^. Second, white matter connectivity (i.e., tractography) can be affected by scanner biases ^12^, which raises doubts about the effectiveness of data harmonization after running whole-brain tractography. Hence, harmonizing the structural connectivity matrices (derived from tractography of unharmonized data) can be inadequate and ineffective in removing scanner effects across dMRI data.

On the other hand, harmonizing dMRI data obtained directly from the scanner has several advantages. As demonstrated in several of our earlier works ^5, 6^, nonlinear and voxel-wise site effects can be removed allowing robust and unbiased tractography estimation across sites. Further, any dMRI microstructural model (e.g. single or multi-tensor, standard model of diffusion, NODDI, etc. ^28–30^) can be used without having to worry about the confounds due to scanner-related biases during the model estimation process. The dMRI community has recognized the value of performing harmonization directly on the scanner data and has organized multiple community challenges to determine the best-performing algorithm as part of the Medical Image Computing and Computer-assisted Intervention (MICCAI) conference ^9, 10^. Across all metrics, our algorithm resulted in the best performance ^9, 10^). The effectiveness of our approach has also been demonstrated through several studies in schizophrenia and other disorders ^6, 7, 31–43^.

Furthermore, the analysis of very large dMRI data sets represents a significant computational burden for many neuroscience researchers and laboratories. Of particular neuroscientific interest is the quantitative hypothesis-driven study of the brain’s major white matter fiber tracts ^44^. To enable the study of the fiber tracts across very large dMRI datasets, it is critical to define anatomical fiber pathways (e.g., the arcuate fasciculus) consistently across all subjects, irrespective of their age, gender, or disease indications. It is also critical to automate the extraction of these fiber pathways and the measurement of their tissue microstructure. Several approaches have demonstrated the advantages of automated extraction of white matter fiber tracts ^45–49^. In this study, we apply our robust multi-fiber tractography method, which allows the measurement of fiber-specific microstructural properties ^50, 51^. We then perform automated tract extraction using a well-established fiber clustering pipeline ^52, 53^ together with a neuroanatomically curated white matter atlas ^45^. We have previously demonstrated that this framework consistently parcellates white matter tracts across the lifespan ^45^ with high test-retest reproducibility ^54^.

The aim of this study is to create a database of harmonized dMRI data and tract-specific microstructure measures to enable novel scientific investigations across thousands of subjects from the ABCD study. To produce this novel database, which includes data from quality controlled 9345 subjects of the over 10,000 subjects, we applied our advanced dMRI harmonization, tractography, and white matter parcellation computational pipeline. First, our robust dMRI data harmonization algorithm was applied to remove scanner-specific biases from the multi-site dMRI data. Harmonizing the raw data acquired directly from the scanner in this way enables laboratories to then perform any desired analyses to test their neuroscientific hypotheses of interest. Second, robust multi-fiber tractography was computed in the whole brain of all 9345 subjects for highly consistent tracing of white matter connections. Third, 73 subject-specific anatomical white matter tracts were extracted from the tractography of each subject in an automated fashion, and their fiber-specific microstructural properties were quantified. The release of this fiber tract microstructure data will enable laboratories to directly test neuroscientific hypotheses of interest without needing to perform computationally intensive processing. This paper includes several experiments that demonstrate the efficacy of dMRI data harmonization and the success of white matter tract identification on the ABCD dataset. Overall, the release of the proposed database represents an unparalleled opportunity for researchers to investigate neurodevelopmental brain changes that were previously challenging to investigate using unharmonized data or smaller sample sizes. The database includes harmonized dMRI data, tractography, extracted white matter tracts, and microstructure measures from the ABCD study. The database is now available through the NIMH Data Archive repository, allowing open access to the neuroimaging and neuroscience community.

## 2) Methods

### 2.1 The ABCD study database

The ABCD study has collected an extensive dataset of 11,878 participants across 21 sites with a baseline age of 8-11 years to gain a better understanding of neurodevelopment between childhood and early adulthood (Figure 1a). The dataset provides a longitudinally collected wealth of measured attributes, such as neuroimaging, cognitive, biospecimen, behavioral, youth self-report and parent self-report metrics, and environmental data. The ABCD dataset can be publicly accessed by researchers through a data use agreement with the NDA. Notably, the neuroimaging data includes dMRI, structural MRI (both T1- and T2-weighted), and functional MRI data, all of which were collected every two years for almost all participants (48% of whom are female). The participants were scanned using Siemens, Philips, and GE scanners with similar acquisition parameters across 21 sites.

**Figure 1.**
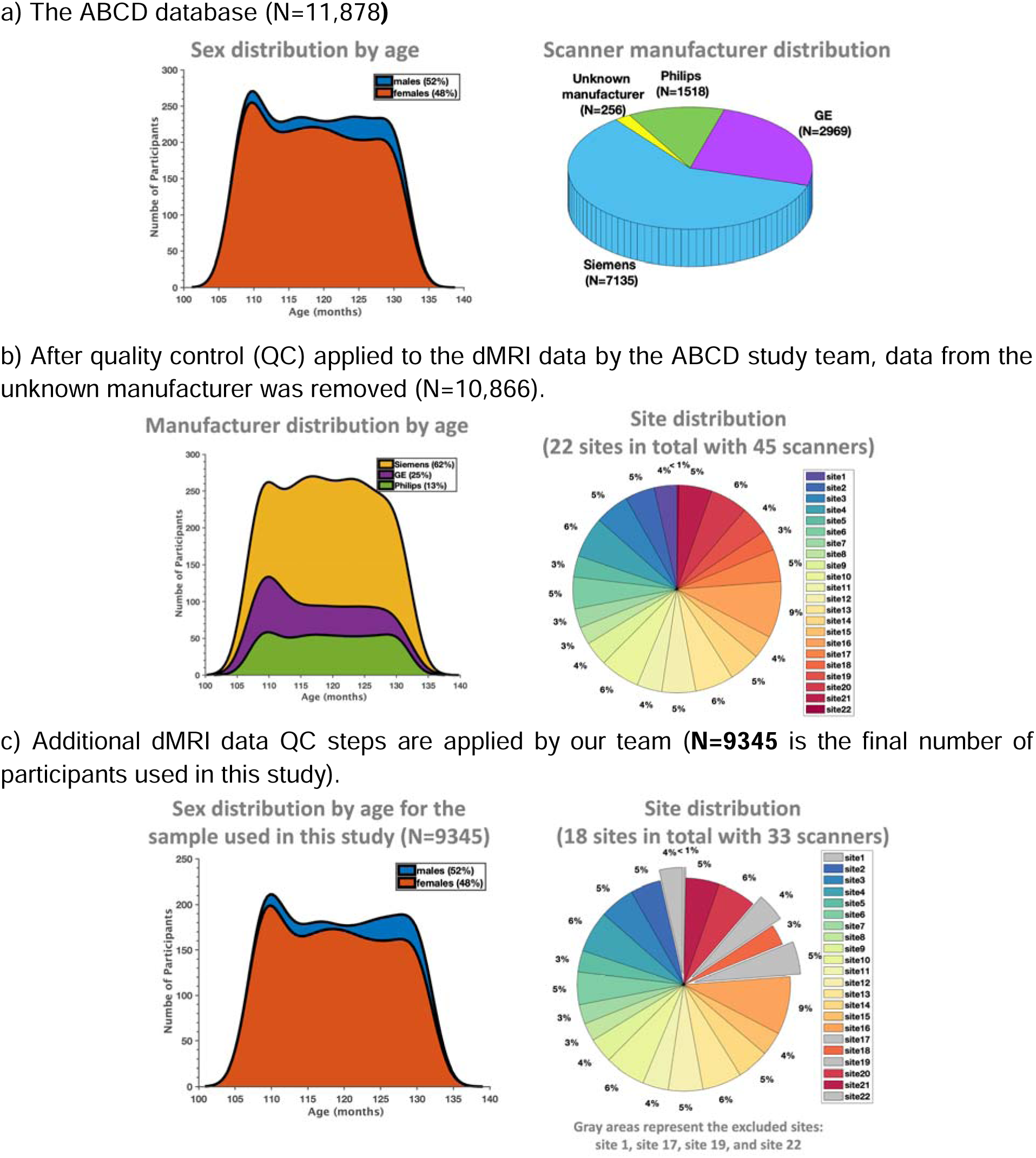
The ABCD study database: (a) The original database before quality control (QC) procedures were applied to the dMRI data (N=11,878); (b) After QC procedures are applied to the dMRI data by the ABCD study team (N=10,866 for the quality passed dMRI scans); (c) Additional QC steps are applied to the dMRI data in this study, where 4 sites with 12 scanners are removed from the sample (N=9345 for the final dMRI scans used in the subsequent processing steps in this study).

The acquired MRI data was minimally preprocessed by the ABCD study Data Analysis, Informatics, and Resource Center group. It is important to note that the ABCD study extensively screened participants for their history of psychiatric disorders, neurological problems, traumatic brain injury (TBI), seizure, low gestational weights, and more. Participants who did not meet the criteria for MRI scanning were excluded from the study, such as those with a history of severe TBI, abnormal MRI scans, or seizures. Detailed information about data recruitment, acquisition, and MRI data preprocessing steps can be found in ^3^ and ^55^. We note that the preprocessing steps of the ABCD study did not include harmonization or tractography analysis of the dMRI data. Therefore, this study is focused on processing multi-site dMRI data from the baseline session using our harmonization and connectivity frameworks, as well as making the processed data of the ABCD study available to the NDA community.

### 2.2 The details on dMRI data acquisition and minimal preprocessing

DMRI scans were acquired at 1.7×1.7×1.7 mm^3^ resolution using multiband EPI with a slice acceleration factor of 3 ^2^. Siemens and GE scanners collected dMRI data using a single acquisition with 96 diffusion gradient directions, seven b=0 volumes, and four b-values (6 directions with b=500 s/mm^2^, 15 directions with b=1000 s/mm^2^, 15 directions with b=2000 s/mm^2^, and 60 directions with b=3000 s/mm^2^). The Philips scanner used two acquisitions to collect the dMRI data. Philips scan 1 parameters were: 3 directions with b=0 s/mm^2^, 4 directions with b=500 s/mm^2^, 7 directions with b=1000 s/mm^2^, 8 directions with b=2000 s/mm^2^ and 29 directions with b=3000 s/mm^2^. Philips scan 2 parameters were: 4 directions with b=0 s/mm^2^, 2 directions with b=500 s/mm^2^, 8 directions with b=1000 s/mm^2^, 7 directions with b=2000 s/mm^2^ and 30 directions with b=3000s/mm^2^ (Table 1). Minimal preprocessing steps were consistently applied to the dMRI data of each study site by the ABCD study, which included: eddy and motion correction, b0 inhomogeneity correction, gradient unwarp, and resampling to isotropic resolution (1.7 mm^3^) ^3,55^. While these preprocessing steps are well established and known to improve data quality, they do not correct for biases due to multiple acquisition sites and scanners, which are addressed as part of this study.

**Table 1.** Data about scanner manufacturers, including the number of subjects scanned by the Siemens/GE/Philips scanners, and the acquisition information. We note that 256 subjects, who do not have scanner information available, are marked as unknown in this table.

| Scanners | Number of subjects | B-values and the number of gradients/volumes |
| --- | --- | --- |
| Siemens | 7135 | *The full protocol consists of 6 b=500, 15 b=1000, 15 b=2000, and 60 b=3000 volumes.* |
| GE | 2969 | *The full protocol consists of 6 b=500, 15 b=1000, 15 b=2000, and 60 b=3000 volumes.* |
| Philips | 1518 | *Includes two acquisitions: 1) 4 b=500, 7 b=1000, 8 b=2000 and 30 b=3000 volumes; 2) 2 b=500, 8 b=1000, 7 b=2000 and 30 b=3000 volumes.* |
| Unknown | 256 | - |
| **Total** | **11,878** |  |

### 2.2 Scanner-Software Upgrades in each ABCD Study Site

Several studies have revealed that even slight modification to the scanner software can result in bias in the acquired dMRI data, similar to a “site-effect,’ even when the acquisition protocol remains unchanged ^13, 56^. However, it is often difficult to predict the bias in the data due to such software updates. Therefore, scanner software (version) updates should ideally be avoided entirely during the course of a study to avoid introducing bias in the dMRI data. However, this is often not feasible in practice. Similar to many large-scale multi-site neuroimaging studies, several scanners within the ABCD study underwent software upgrades during the acquisition of dMRI data. Therefore, we treated each upgrade as an independent scanner. Hence, after counting each software upgrade as a separate “scanner,” the ABCD study included 45 total scanners. Supplementary Table 1 provides details about the scanners and related upgrades.

### 2.3 DMRI Data Quality Control

The ABCD study applied several automated quality control (QC) procedures, such as head motion statistics and the detection of signal drops to remove low quality dMRI scans ^3,55^. The trained technicians at the sites of the ABCD study also did manual checks by reviewing the dMRI scans for poor image quality and various imaging artifacts. The scans that did not pass the QC checks were marked as unacceptable data. Refer to Figure 1b for the distribution of the subjects (per site) that passed the required dMRI data quality.

Additionally, we applied our own automated and manual QC steps to ensure the data was of good quality for harmonization and connectivity analysis (i.e., whole brain tractography and white matter parcellation). This involved using the average whole-brain FA as well as regional FA to screen for outliers, as described in our previous study ^6^. We also conducted manual QC checks on a randomly selected subset of subjects from each scanner. Our findings showed that the dMRI data from most GE and Siemens sites had good quality and was suitable for harmonization and tractography analysis. However, the minimally processed dMRI data from the Philips scanners were found to have certain artifacts, such as ringing and motion artifacts as well as excessive smoothing (potentially during pre-processing steps). Due to these factors, data from the Philips scanners that included 3 sites with 12 scanners used to scan 1518 subjects were excluded from the current study. Our final sample consisted of dMRI data from 9345 participants from Siemens and GE scanners, which were obtained from 18 sites and 33 scanners. Refer to Figure 1c for the distribution of subjects and sites used in this work (i.e., for harmonization and tractography analysis). Supplementary Table 2 provides more details about the sites and scanners used in this work.

### 2.4 Overview of the proposed pipeline to process the dMRI Data (N=9345)

This section describes the major computational steps used to create a database of harmonized and processed dMRI data derived from the ABCD study (see Figure 2). First, brain masks were generated to isolate only the brain tissue and exclude the skull. This was crucial to focus solely on the brain region, making it an indispensable prerequisite for many processing pipelines in neuroimaging studies including ours. To this end, we applied our deep learning-based brain masking tool that robustly extracted the brain from the field-of-view volume in dMRI data of 9345 subjects, as shown in Figure A.1. The details of this method and performance evaluation are described under Appendix A. We then applied the following computational steps respectively: a) dMRI data harmonization, b) whole-brain tractography, c) subject-specific white matter parcellation, d) extracting dMRI measures.

**Figure 2.**
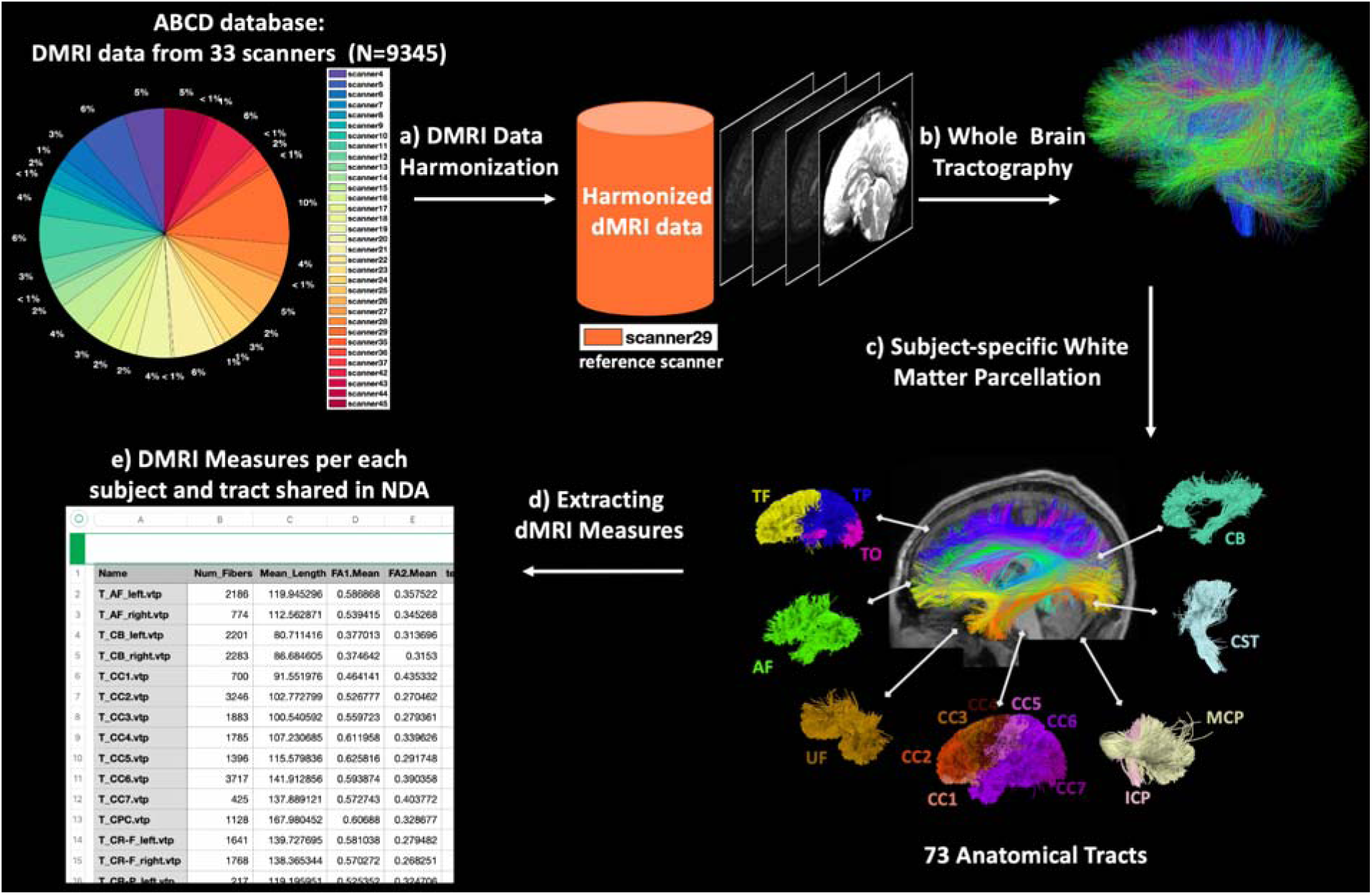
Pipeline overview. (a) DMRI data are harmonized to the reference scanner (scanner 29) to remove possible site/scanner-specific effects. (b) Whole-brain tractography was computed for each subject using the UKF tractography algorithm. (c) WM parcellation was performed using the WMA package in conjunction with an anatomical white matter atlas, resulting in parcellation of 73 anatomically well-defined tracts. (d) Multiple dMRI measures were extracted for each parcellated tract.

#### (a) DMRI Data Harmonization

We applied our retrospective harmonization algorithm to remove the scanner-related differences across 33 scanners. Our harmonization approach is based on rotation invariant spherical harmonics (RISH) features and works at the signal level to remove the scanner-related biases across datasets while accounting for non-linearities in the dMRI data (which can vary by region and tissue).

The harmonization algorithm proceeds by selecting one scanner as the reference and with all other scanners harmonized to this reference site. Scanner 29 from site 16 (a Siemens Prisma scanner) was selected as the reference due to its large sample size (N=959) and high data quality. The dMRI data from the remaining 32 scanners were harmonized to this reference scanner (for a full list, see Supp. Table 2). While scanner 29 was chosen as the reference in this study, we have previously shown that choosing a different scanner as the reference does not affect the performance of harmonization ^6^.

**Table 2.** A total of 73 anatomical fiber tracts are identified for each subject, including 24 association tracts, 8 cerebellar tracts, 7 commissural tracts, 18 projection tracts, and 16 categories of short and medium range superficial fiber clusters organized according to the brain lobes that they connect.

| Tract category (number of tracts) | Tract name (L - left; R - right; C - commissural) |
| --- | --- |
| **Association tracts (24)** | arcuate fasciculus (AF) – LR<br>cingulum bundle (CB) – LR<br>external capsule (EC) – LR<br>extreme capsule (EmC) – LR<br>inferior longitudinal fasciculus (ILF) – LR<br>inferior occipito-frontal fasciculus (IoFF) – LR<br>middle longitudinal fasciculus (MdLF) – LR<br>posterior limb of internal capsule (PLIC) – LR<br>superior longitudinal fasciculus I (SLF I) – LR<br>superior longitudinal fasciculus II (SLF II) – LR<br>superior longitudinal fasciculus II (SLF III) – LR<br>uncinate fasciculus (UF) – LR |
| **Cerebellar tracts (8)** | cortico-ponto-cerebellar (CPC) – C<br>inferior cerebellar peduncle (ICP) – LR<br>middle cerebellar peduncle (MCP) – C<br>intracerebellar input and Purkinje tract (CBLM-I&P) – LR<br>intracerebellar parallel tract (CBLM-PaT) – LR |
| **Commissural tracts (7)** | corpus callosum 1 (CC 1) – C<br>corpus callosum 2 (CC 2) – C<br>corpus callosum 3 (CC 3) – C<br>corpus callosum 4 (CC 4) – C<br>corpus callosum 5 (CC 5) – C<br>corpus callosum 6 (CC 6) – C<br>corpus callosum 7 (CC 7) – C |
| **Projection tracts (18)** | corticospinal tract (CST) – LR<br>corona-radiata-frontal (CR-F) – LR<br>corona-radiata-parietal (CR-P) – LR |
|  | striato-frontal (SF) – LR<br>striato-occipital (SO) – LR<br>striato-parietal (SP) – LR<br>thalamo-frontal (TF) – LR<br>thalamo-occipital (TO) – LR<br>thalamo-parietal (TP) – LR |
| **Superficial tracts (16)** | superficial-frontal (Sup-F) – LR<br>superficial-frontal-parietal (Sup-FP) – LR<br>superficial-occipital (Sup-O) – LR<br>superficial-occipital-temporal (Sup-OT) – LR<br>superficial-parietal (Sup-P) – LR<br>superficial-parietal-occipital (Sup-PO) – LR<br>superficial-parietal-temporal (Sup-PT) – LR<br>superficial-temporal (Sup-T) – LR |

Harmonization of the dMRI data is carried out in two steps. The first step, referred to as *<u>template creation</u>*, involves matching approximately 35 subjects from each scanner with subjects from the reference scanner based on age, sex, and IQ. This matching process is conducted at a group level to eliminate differences between scanners while preserving inter-subject biological variability. To ensure we capture only scanner-related effects, subjects with a history of epilepsy, stroke, tumor, hemorrhage, or psychiatric disorders were not included in the template creation step. The full exclusion criteria can be found in the supplement. The matched set of 35 subjects is then used to learn a nonlinear mapping between dMRI data from different scanners. This inter-scanner mapping is voxel-specific and uses the fiber-orientation-independent RISH feature representation to learn a specific mapping for each RISH feature. These RISH features capture different microstructural tissue properties that are independent of the orientation ^57^ and provide the ability to reconstruct the harmonized dMRI signal. In the second step, *<u>harmonizing dMRI data</u>*, the learned mappings (from the template creation step) across RISH features were applied to the all dMRI data of the corresponding scanner to be harmonized with the reference (including the excluded subjects in the template creation step). Methodological details of these steps can be found in ^6^.

For harmonizing this large-scale dataset (N = 9345), we used the AWS EC2 platform. Specifically, r5d.4xlarge EC2 instances (128GiB of memory, 16 vCPUs) were used in template generation, where it took ∼48 hours to generate each template. We then used r5d.large EC2 instances (16GiB of memory, 2 vCPUs) in the harmonization step, where harmonization of each subject took ∼15 mins. The whole pipeline was tailored for parallel execution in 32 instances. In total, it took about 3872 CPU hours (∼162 CPU days) to complete the processing of all dMRI data (N=9345) in this step.

#### (b) Whole Brain Tractography

*Tractography* was performed using our well-validated multi-tensor unscented Kalman filter (UKF) fiber tracking algorithm ^50, 51, 58, 59^ on the b=3000 s/mm^2^ shell. The two-tensor model adopted in the UKF tractography algorithm is able to depict crossing fibers, which are prevalent in white matter tracts ^60, 61^. In this way, the first tensor is associated with the fiber that is being traced and enables quantification of fiber-specific microstructural properties, while the second tensor models the fibers that cross through the fiber. Moreover, the two-tensor model allows UKF tractography to consistently trace fibers across various populations ^45^ as well as disease states such as tumors ^62–64^. We successfully applied UKF tractography on 9345 subjects from the harmonized ABCD dataset. For each harmonized dMRI data, ∼450k fiber streamlines were generated to create a whole brain tractography dataset.

The UKF tractography was run on the AWS EC2 platform. Specifically, r5d.large EC2 instances (16GiB of memory, 2 vCPUs) were used, where tractography took ∼4.5 hours per subject. A total of ∼500 instances were used, with each instance running tractography for about 20 subjects. In total, it took about ∼42,052 CPU hours/∼1752 CPU days to complete whole brain tractography.

#### (c) Subject-specific White Matter Parcellation

Tractography parcellation was performed using our WhiteMatterAnalysis (WMA) ^65^ fiber clustering pipeline ^52, 66^ in conjunction with our anatomical white matter atlas ^45^. This pipeline produces a fine-scale whole-brain tractography parcellation into 800 fiber clusters and a coarse-scale anatomical tract parcellation, including 57 major deep WM tracts and 16 categories of superficial fiber clusters organized according to the brain lobes they connect. Refer to Table 2 for a list of anatomical tracts. Figure 2 gives a visualization of example tracts from 10 randomly selected subjects from 10 different scanners.

We used the AWS r5d.large EC2 instances (16GiB of memory, 2 vCPUs) for the computation, where it took ∼1.6 hours per subject. A total of ∼500 instances were used, with each instance running white matter parcellation for about 20 subjects. In total, it took about ∼14,952 CPU hours/∼623 CPU days to complete the white matter parcellation.

#### (d) Extracting DMRI Measures

Multiple dMRI measures were extracted from each parcellated cluster and anatomical tract using SlicerDMRI ^67, 68^ for all subjects. These include the widely used dMRI measures: FA, mean diffusivity (MD), axial diffusivity (AD), radial diffusivity (RD), number of streamlines (NoS), number of streamline points (NoF) and streamline length. The tensors associated at each streamline location were also saved in the corresponding VTK file which can be accessed from the NDA. These extracted dMRI measures are available via the NDA repository. Additional measures of interest can also be computed from the tensors by downloading the appropriate VTK file corresponding to a particular anatomical tract or fiber cluster.

## 3) Data Records

### 3.1 Sharing the dMRI measures and processed dMRI data in NDA

The generated imaging files and corresponding dMRI measures for each tract are shared in NDA through two separate submissions.

First, a total of 804 different derived measures, including FA, MD, NoS, etc. for each of the 73 white matter fiber bundles (bilateral, commissural, superficial, and cerebellar) were uploaded on the NDA for 9345 subjects. Refer to Table 3 for the list of shared dMRI measures. These measures will be made available by the NDA during their next data release.

**Table 3.**
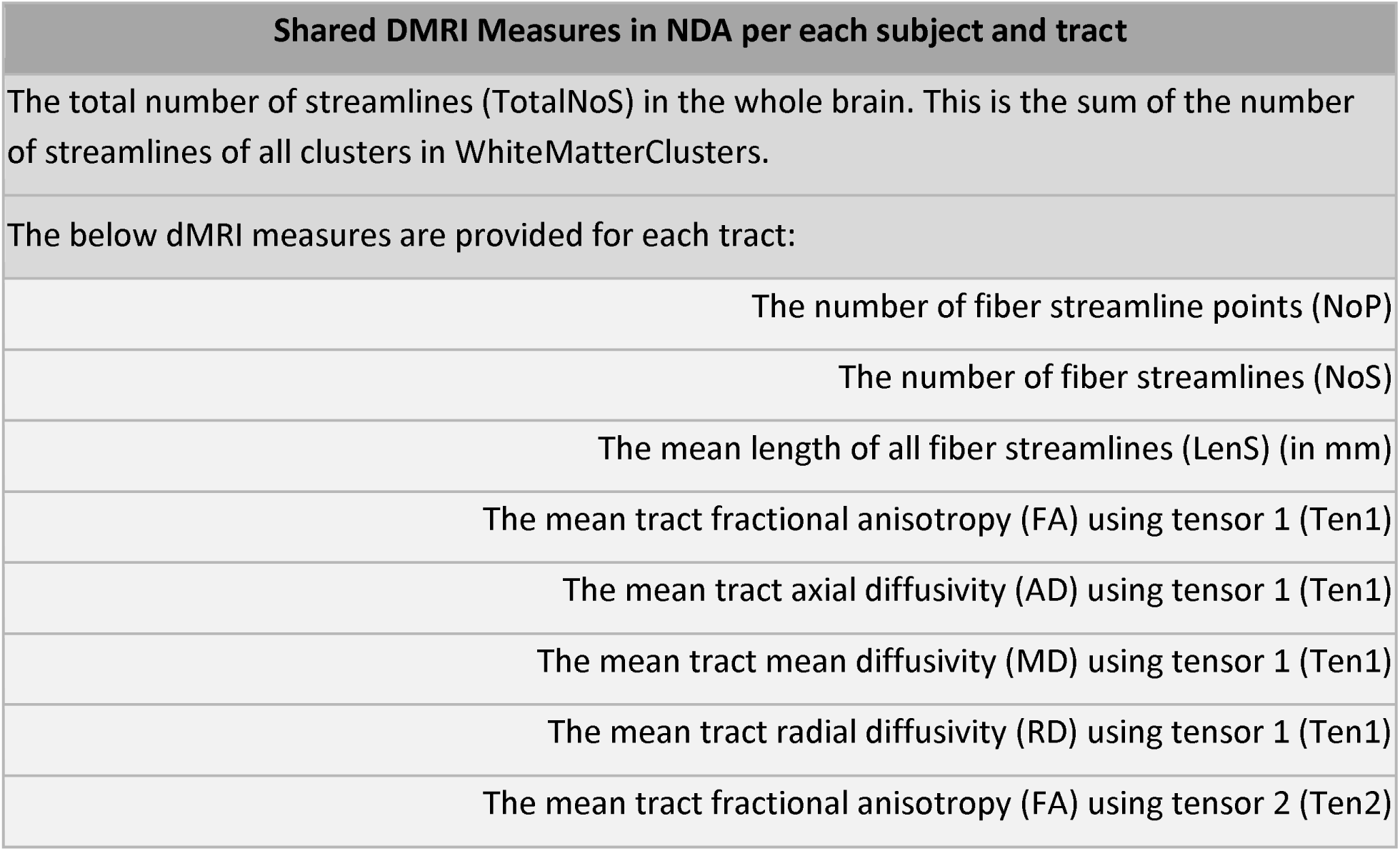
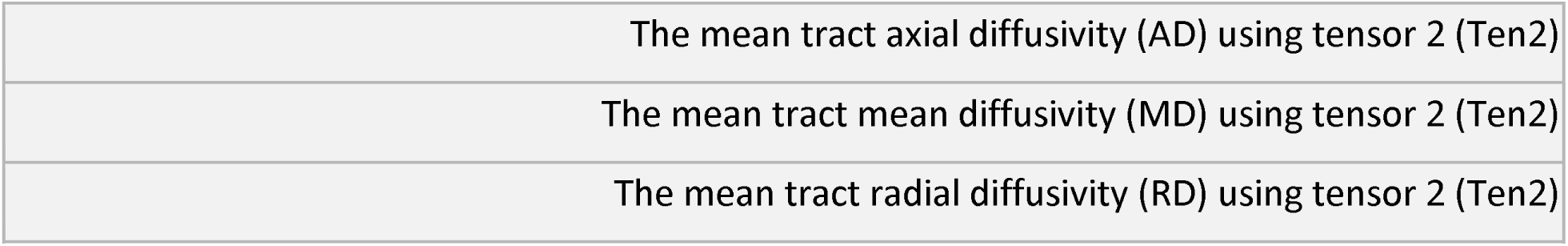
A total of 804 different derived measures, including FA, MD, NoS, etc. for each of the 73 white matter fiber bundles (bilateral, commissural, superficial and cerebellar) were uploaded on the NDA for 9345 subjects.

Second, we shared the harmonized and processed imaging data, including the harmonized dMRI data, brain masks, whole brain tractography, anatomical fiber tracts, and white matter clusters with the NDA community. For the anatomical fiber tracts, we have provided both full-size VTK files for comprehensive visualization and quantitative analysis, as well as downsampled VTK files for quicker download and lower-resolution tract visualization. The full-size VTK files for white matter clusters have also been shared. It should be noted that the shared VTK files contain tensors estimated at each point, allowing for the computation of additional measures if necessary.

### 3.2 Access request in NDA

The standard process for initiating a Data Access Request (DAR) for shared data on the NDA involves selecting a Permission Group from the NDA Data Permissions Dashboard and providing a Research Data Use Statement, contact information for all recipients who will access the data, and the name of a Signing Official (SO) at the lead recipient’s research institution. The Data Use Certification (DUC) PDF is downloaded and signed by the investigators and the SO and then uploaded to the appropriate “active request’ on the NDA Permissions Dashboard. NDA staff reviews the DUC for completeness and sends the request to the Data Access Committee (DAC), which makes decisions based on research subject protection and adherence to data use limitations consented to by the research subjects. Approved recipients gain access to the shared data for one year, and denied recipients may submit a new request that addresses the DAC’s reason for denial. The details for this process can be found on the NDA’s website: https://nda.nih.gov/nda/access-data-info.html

Access to our harmonized dMRI data and measures requires only permission from the standard “NIMH Data Archive’ permission group. This permission group enables standard access to phenotypic, imaging, and genomic data, as well as supporting documentation for NIH-funded grants for a period of one year. Renewal requests must be submitted at the end of each year to maintain access.

### 3.3 Downloading the harmonized dMRI data and measures

Once access to NDA has been granted, users can utilize the NDA webpage and NDA Query Tool to search our project titled “Harmonizing multi-site diffusion MRI acquisitions for neuroscientific analysis across ages and brain disorders”. Please find below the step-by-step instructions to access and download the datasets of harmonized ABCD dMRI data and measures from the NDA:

1. Launch a web browser and navigate to the NDA website at https://nda.nih.gov/. Proceed to log in to the system using the provided login credentials.
2. Click on the “Get Data’ option on the website’s primary menu, or utilize the link https://nda.nih.gov/general-query.html?q=query=collections%20~and~%20orderBy=id%20~and~%20orderDirection=Ascending to access the “Get Data’ page directly.
3. Employ the NDA Query Tool and input one of the following search criteria into the “Text Search’ box to locate our project:

- Enter the full project title: “Harmonizing multi-site diffusion MRI acquisitions for neuroscientific analysis across ages and brain disorders.”
- Alternatively, search using keywords such as “Harmonize AND ABCD.”
4. Once our project is located, select it by clicking on the “Add to Workspace’ button to add the dataset to your personal workspace.
5. Proceed to your “Workspace’ and click on it to submit the dataset to the “Filter Cart.’ The cart may take a few moments to update.
6. Within the “Filter Cart,’ click on “Create Data Package/Add Data to Study’ to access the Data Packaging Page.
7. On the Data Packaging Page, click on “Create Data Package.’ Assign a unique name to the package, and ensure that “include associated data files’ is selected to download the imaging data. To monitor the status of your package, access “Data Packages’ on your user profile, and periodically refresh the webpage until the package is ready for download.
8. Finally, install the “NDA Download Manager’ from the following link: https://nda.nih.gov/nda/nda-tools.html#download-manager. Use this tool to download your data package once the package is ready to download.

Please note that this interface might be updated by the NDA in the next few years, in which case, we would ask the user to refer to their updated manuals for accessing the data.

## 4) Technical Validation

### 4.1 Descriptions of the post-harmonization experiments on the ABCD dataset

The baseline dMRI data from 9345 subjects of the ABCD study underwent harmonization, which brought the dMRI data of all 32 scanners from 18 different sites in line with the reference dataset (dataset #29 from site 16). In this section, we describe two experiments to demonstrate effectiveness of the harmonization algorithm to remove scanner effects from the baseline dMRI data in the ABCD study (Figure 4).

**Figure 3.**
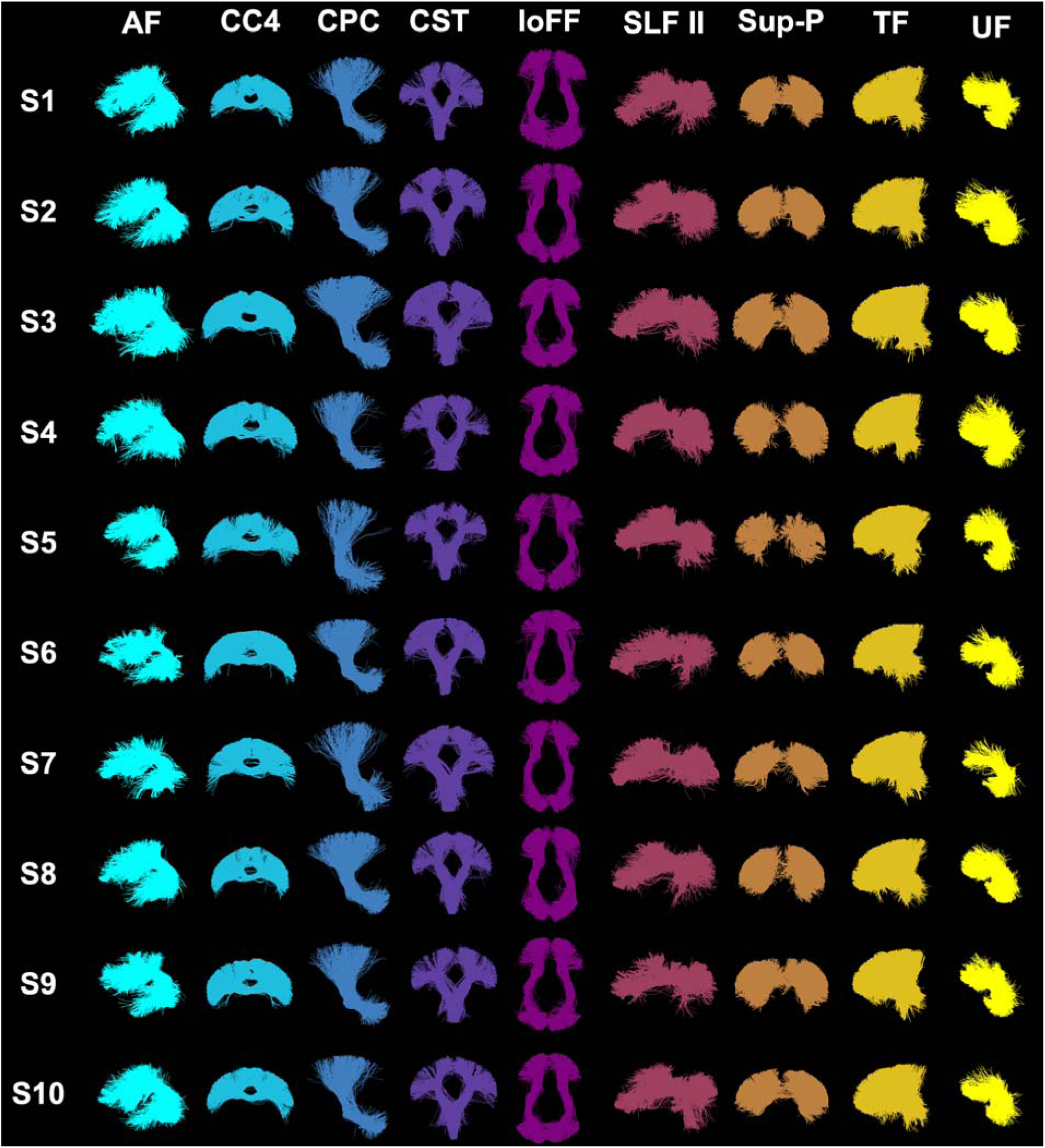
Visualization of example anatomical tracts. 10 randomly selected subjects (from 10 different sites) are used. In general, the parcellated anatomical tracts are visually highly similar across the different subjects. Refer to Table 2 for the definition of the acronyms for tract names.

**Figure 4.**
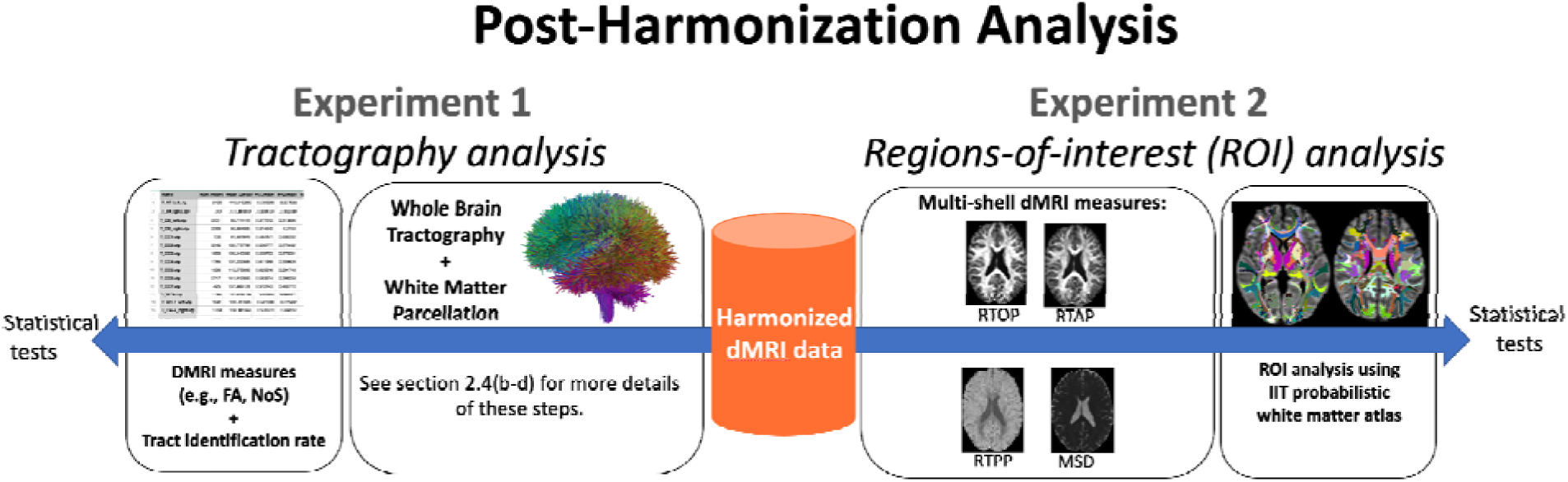
Overview of the experiments that demonstrate the effectiveness of harmonization. Experiment 1 evaluated the impact of harmonization on dMRI measures, such as FA, and tract identification rate, obtained through tractography analysis and white matter parcellation. In contrast, Experiment 2 assessed the impact of harmonization on multi-shell dMRI measures, such as RTOP, RTAP, RTPP, and MSD, using white matter ROIs defined by the IIT probabilistic white matter atlas.

#### (a) Experiment 1: Tractography analysis

We first assessed the effects of dMRI harmonization on white matter parcellation (Section 2.4.c) and the extracted dMRI measures (Section 2.4.d) at the tract level. Specifically, for each tract per scanner, we measured the identification rate ^45, 47^ to quantify the quality of the tracts detected before and after harmonization. Here, a tract is considered to be successfully identified if there are at least 10 streamlines inside the tract ^45, 47, 69^, and its identification rate is defined as the percentage of successfully identified tracts across all analyzed subjects. In three randomly selected sites, we repeated this experiment on the original dMRI data to be able to compare the identification rate before and after harmonization. In addition, for each tract, we also computed the mean FA (from tensor 1) across all subjects for each dataset before and after harmonization and compared it to the reference data (to which all datasets were harmonized). We repeated the entire analysis at the cluster level as well to demonstrate the effects of dMRI harmonization on the clusters.

#### (b) Experiment 2: Regions-of-interest (ROI) analysis

We further evaluated the performance of the harmonization on multi-shell dMRI measures. To this end, we computed several multi-shell dMRI measures such as Return-To-Origin Probability (RTOP), Return to Axis Probability (RTAP), Return to the Plane Probability (RTPP), and Mean Squared Displacement (MSD) ^28, 70, 71^ in the original, harmonized and reference dMRI datasets using all b-value shells in the dMRI data. We note that for computational efficiency, we compared these measures across the 35 matched subjects that were used in the template creation step. Averages of the multi-shell dMRI measures were calculated over the whole brain white matter skeleton and 42 white matter regions using the Illinois Institute of Technology’s (IIT) brain ROI atlas in the standard MNI space ^72, 73^. To evaluate the performance of the harmonization, we compared the original and harmonized datasets to the reference dataset using unpaired t-tests.

To further verify the effectiveness of harmonization, we conducted a test using a set of 35 subjects that were not included in the process of creating the template for harmonization. These subjects were selected from the dataset that was being harmonized to the reference data, and were chosen based on their similarity in age, sex, and IQ to the reference data, in order to minimize any biological differences that could affect the results. This test on the unseen dataset was carried out to replicate the ROI analysis results in an independent dataset that was not used during the template creation (i.e., the learning process).

### 4.2 Results of the post-harmonization experiments on the ABCD study

The harmonization demonstrated consistent results across all dMRI measures in two separate experiments. The outcomes of each of these experiments are detailed in the following sections.

#### (a) Experiment 1: Tractography analysis

The impact of harmonization on dMRI measures obtained through tractography analysis was evaluated. The study focused on three randomly selected sites that used different scanners (Siemens Prisma, Siemens Prisma-fit, GE) to investigate the effect of harmonization on white matter parcellation and extracted dMRI measures. The analysis included both cluster- and tract-level evaluations. Figure 5 illustrates the effect of harmonization on white matter parcellation. Harmonization was found to improve the identification rates in both cluster- and tract-level parcellations (see Figure 6 (a) and (c)) for all three scanners). Furthermore, the mean cluster and tract FA values (Figure 6 (b) and (d)) were found to be closer to the reference data after harmonization. Across all 9345 harmonized dMRI data from 21 sites in the ABCD study, high identification rates were observed in both tract- and cluster-level parcellations, at 99.9% and 97.5%, respectively. Figure 6 demonstrates the average FA comparisons for each tract individually. Any significant differences that were observed between the reference and original datasets (p<0.01) were eliminated after harmonization (p>0.1).

**Figure 5.**
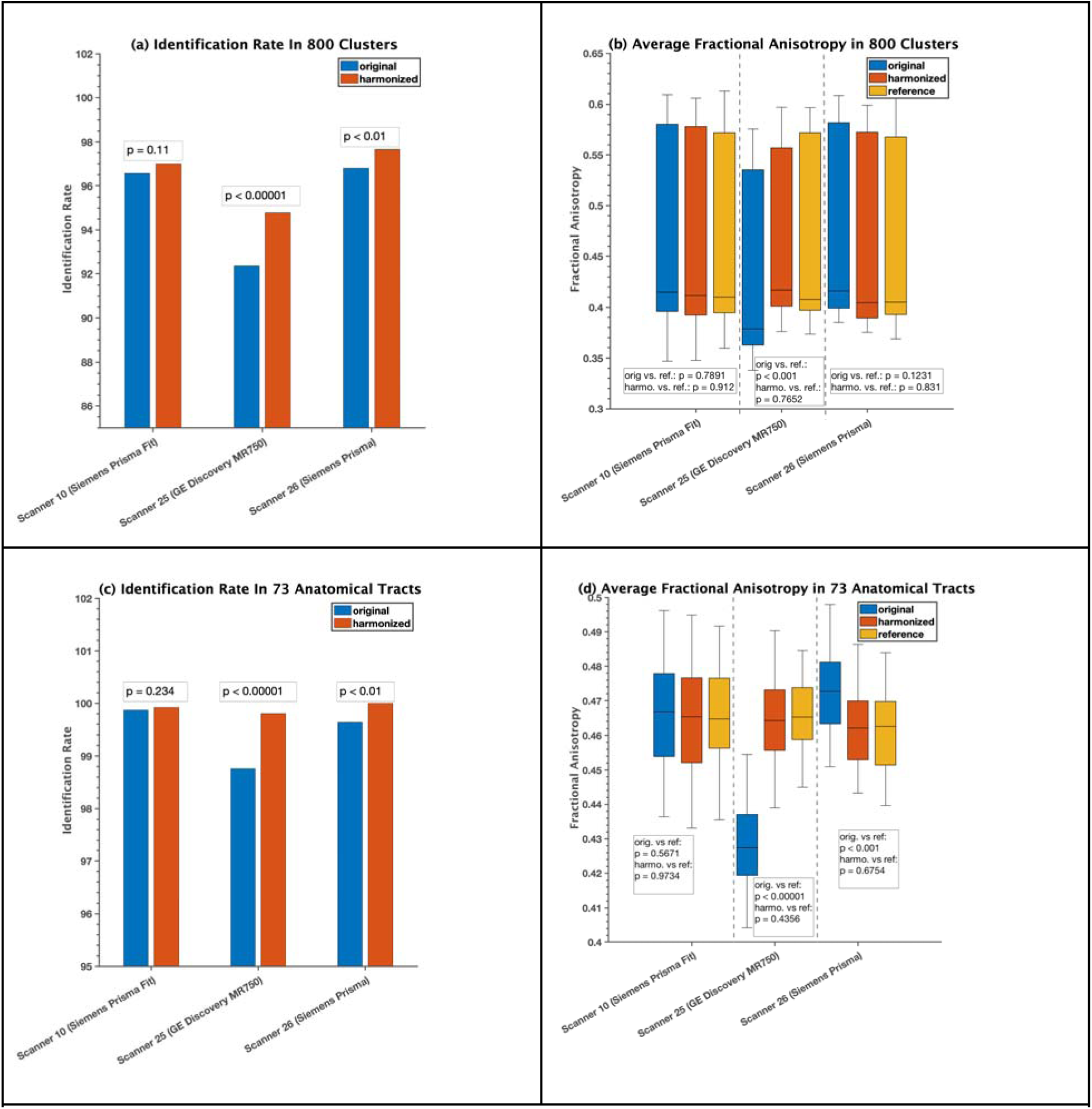
Experiment 1, section 2.6.a, investigated the effects of dMRI harmonization on white matter parcellation in three randomly selected scanners. For this analysis, thirty-five (35) subjects from each scanner, who were part of the learning process of harmonization, were selected for comparison. These subjects were matched in terms of age, sex, and IQ to the reference scanner (scanner 29). The cluster-level analysis is presented in (a) and (b), while the tract-level analysis is shown in (c) and (d). In (a) and (c), a paired t-test was conducted for each scanner to determine whether there was a significant improvement in the identification rate after harmonization. In (b) and (d), two unpaired t-tests were performed for each scanner to compare the average FA between (i) the original and reference scanners and (ii) the harmonized and reference scanners. The resulting p-values were presented on the plots.

**Figure 6.**
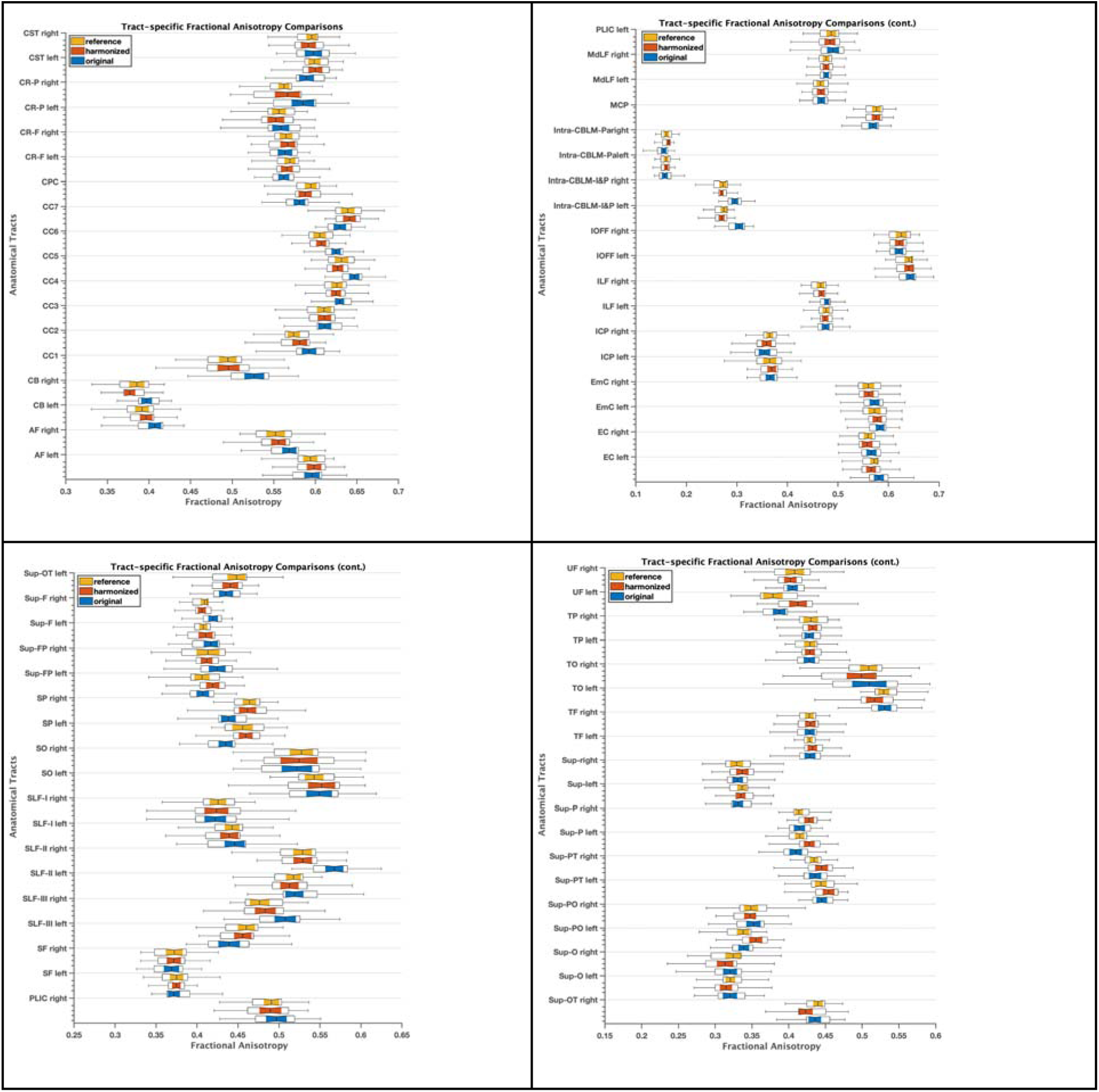
Experiment 1, section 2.6.a, demonstrated the effects of dMRI harmonization on 73 white matter tracts by comparing the average FA of reference, original, and harmonized datasets. Table 2 defines the acronyms for the tract names. For this analysis, 35 subjects from scanner 10 (Siemens Prisma fit) were selected for comparison. These subjects were used as part of the learning process of harmonization and were matched in terms of age, sex, and IQ to the reference scanner (scanner 29). Any significant differences observed prior to harmonization (p<0.01) were eliminated after harmonization (p>0.1).

#### (b) Experiment 2: Regions-of-interest (ROI) analysis

The impact of harmonization on the multi-shell dMRI measures (RTOP, RTAP, RTPP, and MSD) was evaluated. The outcome of this experiment is presented in Figure 7, which demonstrates a comparison of the average multi-shell dMRI measures for different white matter ROIs across eight distinct scanners. The comparison was made between the original data (before harmonization), harmonized data, and reference data. The results indicated significant differences (p<0.01) in the multi-shell dMRI measures between the original and reference datasets. However, after harmonization, these differences were reduced and statistically diminished (p>0.1).

**Figure 7.**
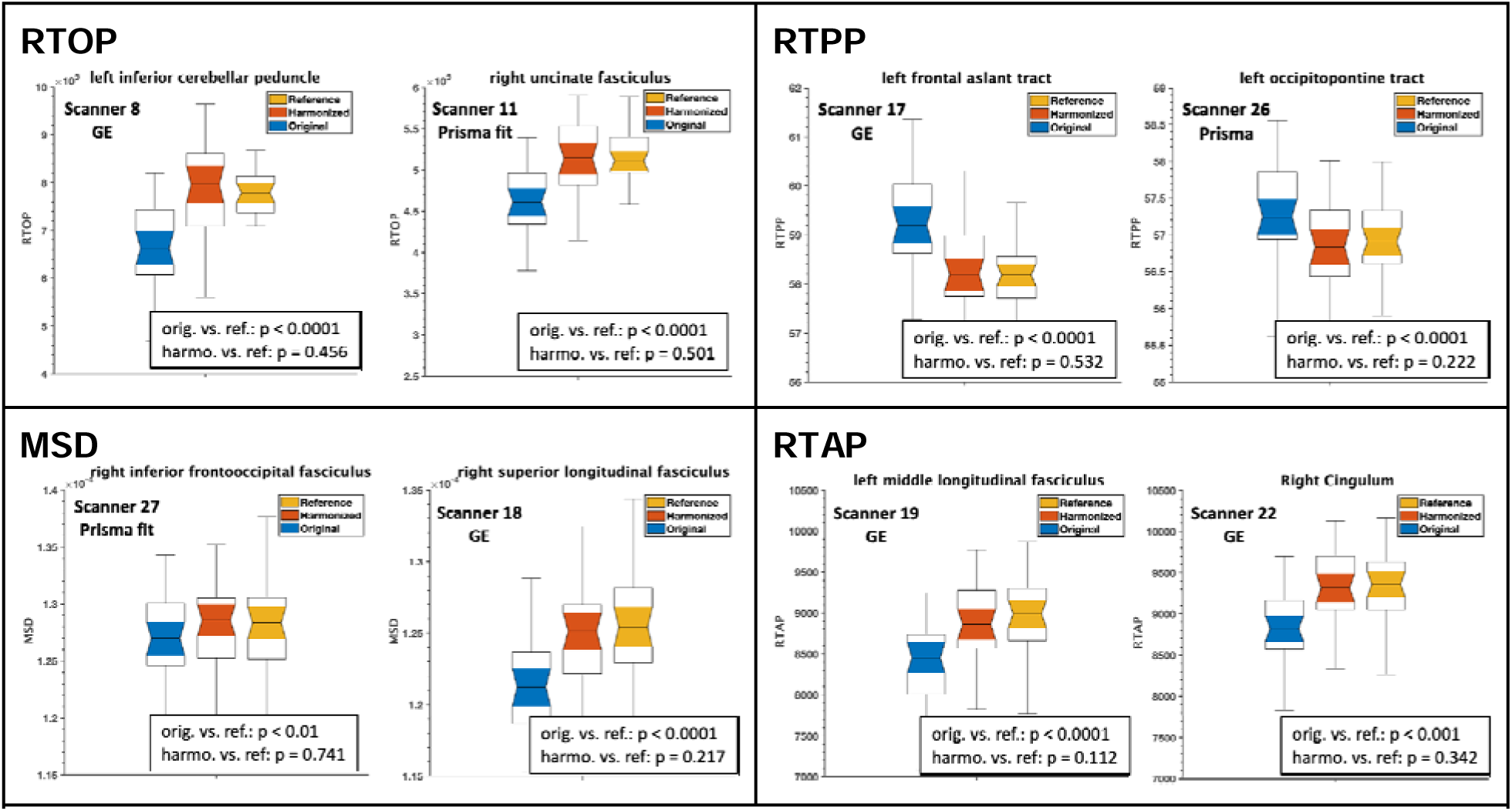
Experiment 2, section 2.6.b, demonstrated the effects of dMRI harmonization on ROI analysis and multi-shell dMRI measures, namely RTOP, RTPP, MSD, RTAP, in eight randomly selected scanners and white matter ROIs. **Thirty-five (35) subjects from each scanner were chosen to create a template for harmonization**, and they were matched in terms of age, sex, and IQ to the reference scanner (scanner 29). Using these subjects from each scanner, two unpaired t-tests were conducted to compare each multi-shell dMRI measure between (i) the original and reference scanners (orig. vs. ref.) and (ii) the harmonized and reference scanners (harmo. vs. ref.). The resulting p-values were presented on the plots.

To further validate the performance of harmonization, we repeated experiment 2 on the dMRI data of new and unseen subjects (i.e., these were not included in the learning process of harmonization/template creation). These subjects were selected from eight scanners and matched with the reference dataset in terms of age, sex, and IQ. The results of the experiment are demonstrated in Figure 8, which compares the average RTOP, RTAP, RTPP, and MSD measures of the several white matter ROIs between the reference, original, and harmonized datasets. Once again, harmonization eliminated any existing scanner-related differences between the original and reference datasets.

**Figure 8.**
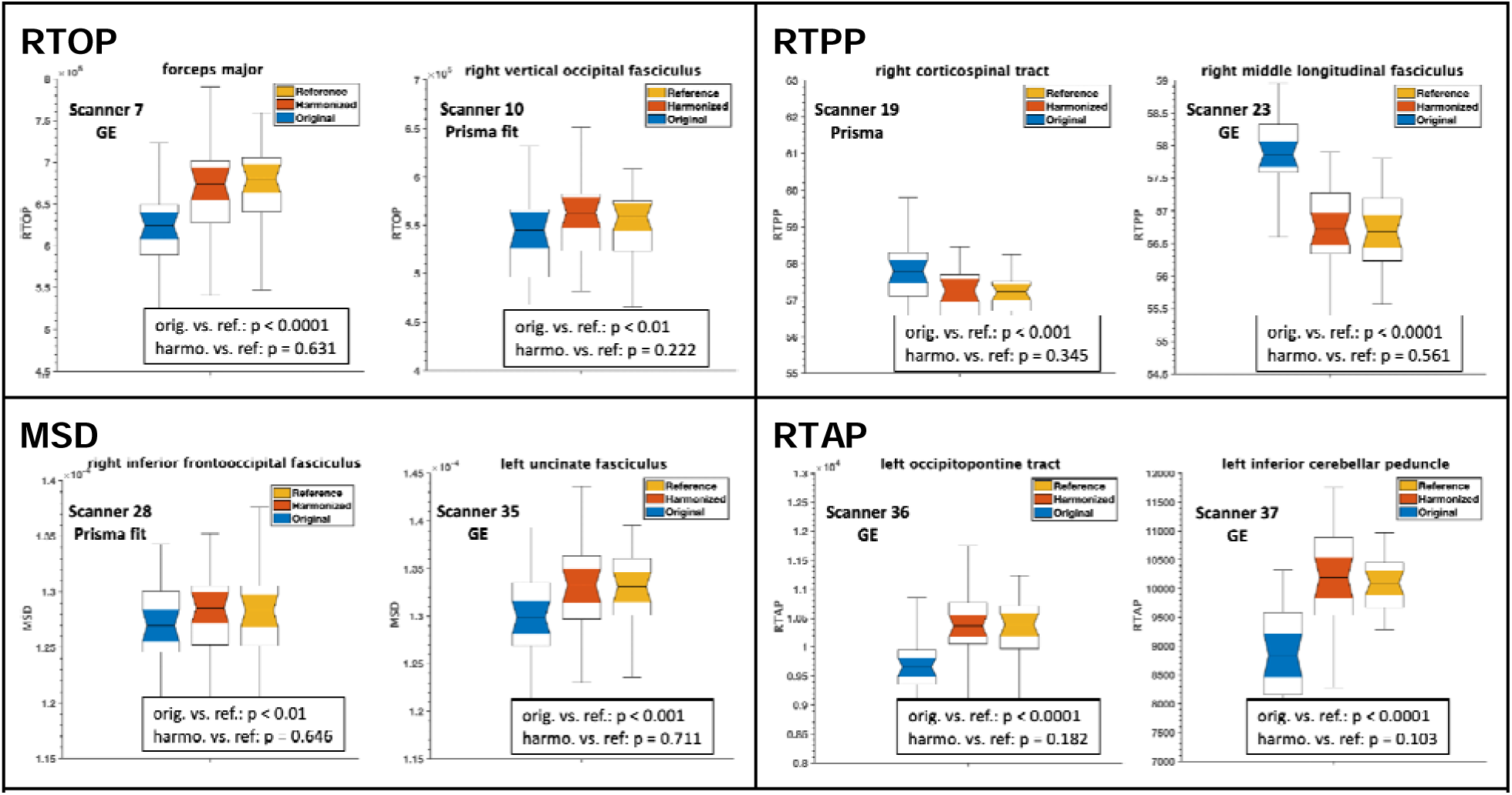
Experiment 2, section 2.6.b, investigated the effects of dMRI harmonization on ROI analysis and multi-shell dMRI measures, namely RTOP, RTPP, MSD, RTAP, in eight randomly selected scanners and white matter ROIs. For this analysis, **an unseen set of thirty-five (35) subjects who were not involved in creating the template** and were still matched in terms of age, sex, and IQ to the reference scanner (scanner 29) were selected. Using these subjects from each scanner, two unpaired t-tests were conducted to compare each multi-shell dMRI measure between (i) the original and reference scanners (orig. vs. ref.) and (ii) the harmonized and reference scanners (harmo. vs. ref.). The resulting p-values were presented on the plots.

## 5) Usage Notes

The purpose of this study was to create a harmonized and processed dMRI database for the ABCD study, which would enable researchers to analyze the entire dMRI data together. The study involved a significant computational undertaking, which required approximately 50,000 CPU hours to successfully harmonize the dMRI data, perform tractography on the entire brain, along with white matter parcellation into 800 clusters per subject. The results of this effort have been made available to the scientific community via the NDA. Researchers can query the project titled “Harmonizing multi-site diffusion MRI acquisitions for neuroscientific analysis across ages and brain disorders’ on the NDA website using the NDA access (see section Data Records for details on how to download the shared data).

The database includes the output of the harmonization algorithm, brain masking and whole brain tractography, as well as white matter tracts, white matter clusters, and derived dMRI-related measures. The measures of the white matter anatomical tracts extracted from the harmonized and processed dMRI data are accessible through the NDA. This includes a total of 808 different derived measures, including FA, MD, NoF, etc., for each of the white matter fiber bundles (73 bilateral, commissural, superficial, and cerebellar) per each subject. Leveraging the harmonized and processed dMRI data will allow for pooled large-scale data analysis as if the data came from the same scanner, which will significantly increase the statistical power of all neuroimaging studies using this comprehensive dataset, the ABCD study.

However, it is important to note that the study has limitations. One primary limitation is that the dMRI scans acquired on the Philips scanner had to be excluded as they did not pass quality checks. Although the excluded scans make up a small portion of the overall dMRI data (less than 13% of all dMRI scans), these subjects can still be distinct enough to affect the statistical significance of studies investigating specific biology- or disease-related characteristics. Additionally, all dMRI data included in this sample comes from only baseline sessions. The ABCD study has been collecting dMRI data for each subject every two years, and future work will focus on harmonizing the dMRI data from following years to provide the ability to analyze the longitudinal dMRI data (e.g., characterize long-term white matter changes).

In conclusion, the harmonized database, which pools together data from multiple sites and scanners without significant scanner bias, will significantly enhance the statistical power of research based on the ABCD study. This will provide the ability to run more advanced statistical and machine learning analyses aimed at uncovering the neurodevelopmental changes in the white matter of adolescents.

## 6) Code Availability

The open access dMRI data processing software used in this study can be accessed on GitHub. The repositories contain the processing pipeline and scripts used for harmonizing and processing the dMRI data, as well as for performing whole brain tractography and white matter parcellation. Researchers and clinicians interested in utilizing these tools for their own research or clinical applications can easily download and customize the software to fit their specific needs.

Additionally, the GitHub repositories include detailed documentation on how to use the software, as well as examples of how to run the scripts on sample data. The repositories also provide information on the dependencies required to run the software, ensuring that researchers have access to all the necessary tools to use the software effectively.

By making the dMRI data processing software openly accessible on GitHub, we hope to encourage further research and clinical applications of the software and facilitate collaboration across the scientific community. Please refer to the following GitHub links for each of the specific dMRI data processing software:

a. Convolutional neural network dMRI brain segmentation: https://github.com/pnlbwh/CNN-Diffusion-MRIBrain-Segmentation
b. DMRI data harmonization: https://github.com/pnlbwh/dMRIharmonization
c. UKF two tensor whole brain tractography: https://github.com/pnlbwh/ukftractography
d. White Matter Analysis: https://github.com/SlicerDMRI/whitematteranalysis
e. SlicerDMRI: http://dmri.slicer.org

## Acknowledgments

We gratefully acknowledge funding provided by the following National Institutes of Health (NIH) grant: R01 MH119222 (PIs: Dr. Yogesh Rathi, Dr. Lauren J. O’Donnell). We also acknowledge funding provided by the Brigham and Women’s Hospital Program for Interdisciplinary Neurosciences through a gift from Lawrence and Tina Rand and the Brain and Behavior Research Foundation NARSAD Young Investigator Award (PI: Dr. Suheyla Cetin-Karayumak).

Data used in the preparation of this article were obtained from the Adolescent Brain Cognitive DevelopmentSM (ABCD) Study (https://abcdstudy.org), held in the NIMH Data Archive (NDA). This is a multisite, longitudinal study designed to recruit more than 10,000 children age 9-10 and follow them over 10 years into early adulthood. The ABCD Study® is supported by the National Institutes of Health and additional federal partners under award numbers U01DA041048, U01DA050989, U01DA051016, U01DA041022, U01DA051018, U01DA051037, U01DA050987, U01DA041174, U01DA041106, U01DA041117, U01DA041028, U01DA041134, U01DA050988, U01DA051039, U01DA041156, U01DA041025, U01DA041120, U01DA051038, U01DA041148, U01DA041093, U01DA041089, U24DA041123, U24DA041147. A full list of supporters is available at https://abcdstudy.org/federal-partners.html. A listing of participating sites and a complete listing of the study investigators can be found at https://abcdstudy.org/consortium_members/. ABCD consortium investigators designed and implemented the study and/or provided data but did not necessarily participate in the analysis or writing of this report. This manuscript reflects the views of the authors and may not reflect the opinions or views of the NIH or ABCD consortium investigators.

## Author Contributions

The study involved multiple authors with distinct roles and responsibilities. SCK was responsible for the data preparation, setting up AWS platforms, running brain masking and harmonization, and drafting the manuscript. FZ was responsible for running the whole brain tractography and white matter analysis pipelines. TB played a role in developing the harmonization software, while LZ contributed to preparing the white matter atlas figures. NM was involved in identifying the white matter tracts, and SP created the low-resolution white matter tracts for visualization. The study was conceived by LJO and YR, who participated in its design and coordination and contributed to drafting the manuscript. All authors carefully reviewed and approved the final manuscript.

## Appendix

### A. Convolutional Neural Network (CNN) DMRI Brain Segmentation.

The CNN architecture in this method is based on the study of ^74^, which was trained to skull-strip the T1-weighted images. In our case, however, we trained the network on baseline (b=0 s/mm^2) images of the dMRI data to separate the brain from non-brain regions in the image. To further improve the segmentation, we modified the CNN architecture by integrating a multi-view aggregation step ^75^. This step combined the results from models trained on 2D slices along three primary axes: coronal, sagittal, and axial. The final brain mask is obtained by combining the probability maps from all three networks.

The brain masking process included a pre-processing step where we registered the b0 images to the standard MNI space along with performing data normalization (scaling the b0 values between 0 and 1) to improve the performance of the model in the training step. The deep learning models were trained using manually curated masks of dMRI data from 1500 subjects collected from 10 different datasets, including 200 subjects from the ABCD study. We employed the Adam optimizer with a learning rate of 1e-3 during the training process, and all networks were run for up to 10 epochs. We implemented the deep learning algorithm in Keras-Tensorflow 2.2.4. Finally, as a post-processing step, we transformed the resulting brain mask from the MNI space back to the subject space. These steps are summarized in Figure A.2.

To demonstrate the effectiveness of our method with respect to existing tools, we compared our deep learning brain masking method with the most commonly used approach, Brain Extraction Tool (BET) ^76^. We evaluated approximately 400 manually corrected masks from the ABCD study with the output of our deep learning network, resulting in a Dice overlap coefficient of 0.99 and a Jaccard index of 0.987. We note that none of these ∼400 subjects were part of the training dataset. In contrast, BET produced lower performance on the same dataset, with a Dice overlap coefficient of 0.957 and a Jaccard index of 0.945.

Finally, we deployed this brain masking tool on the dMRI data of 9345 subjects from the ABCD study using Amazon Web Services (AWS) Elastic Compute Cloud (EC2) g4dn.2xlarge instances (32 GiB CPU memory, NVIDIA T4 GPU with 32 GiB GPU memory, 8 vCPUs), where the entire brain masking process took about 1 min and 30 seconds for each subject. Refer to Figure A.1 for the demonstration of the dMRI data brain masks on 5 randomly picked brains.

**Figure A.1.**
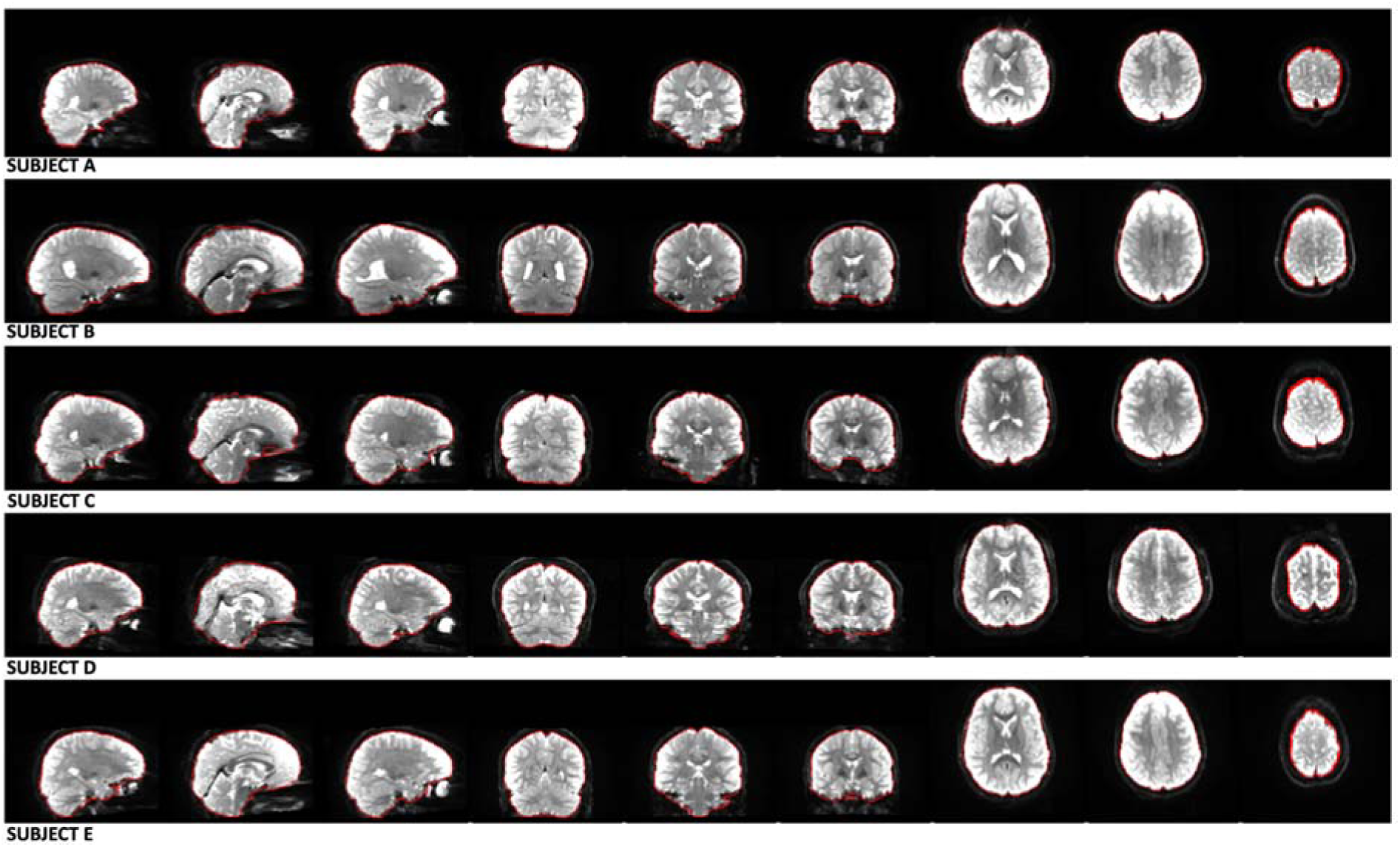
Our deep learning CNN method (https://github.com/pnlbwh/CNN-Diffusion-MRIBrain-Segmentation) was used to perform dMRI Brain Segmentation on the ABCD study’s dMRI data. The results of the segmentation are denoted by the red outlines, which successfully delineate the brains of five randomly selected subjects (Subject A, B, C, D, E) from the ABCD study. We employed fsl’s slicesdir (Jenkinson et al. 2012) for visualization purposes, and each row of the visualization represents a different subject’s brain, while each column displays the various brain slices of that subject.

**Figure A.2.**
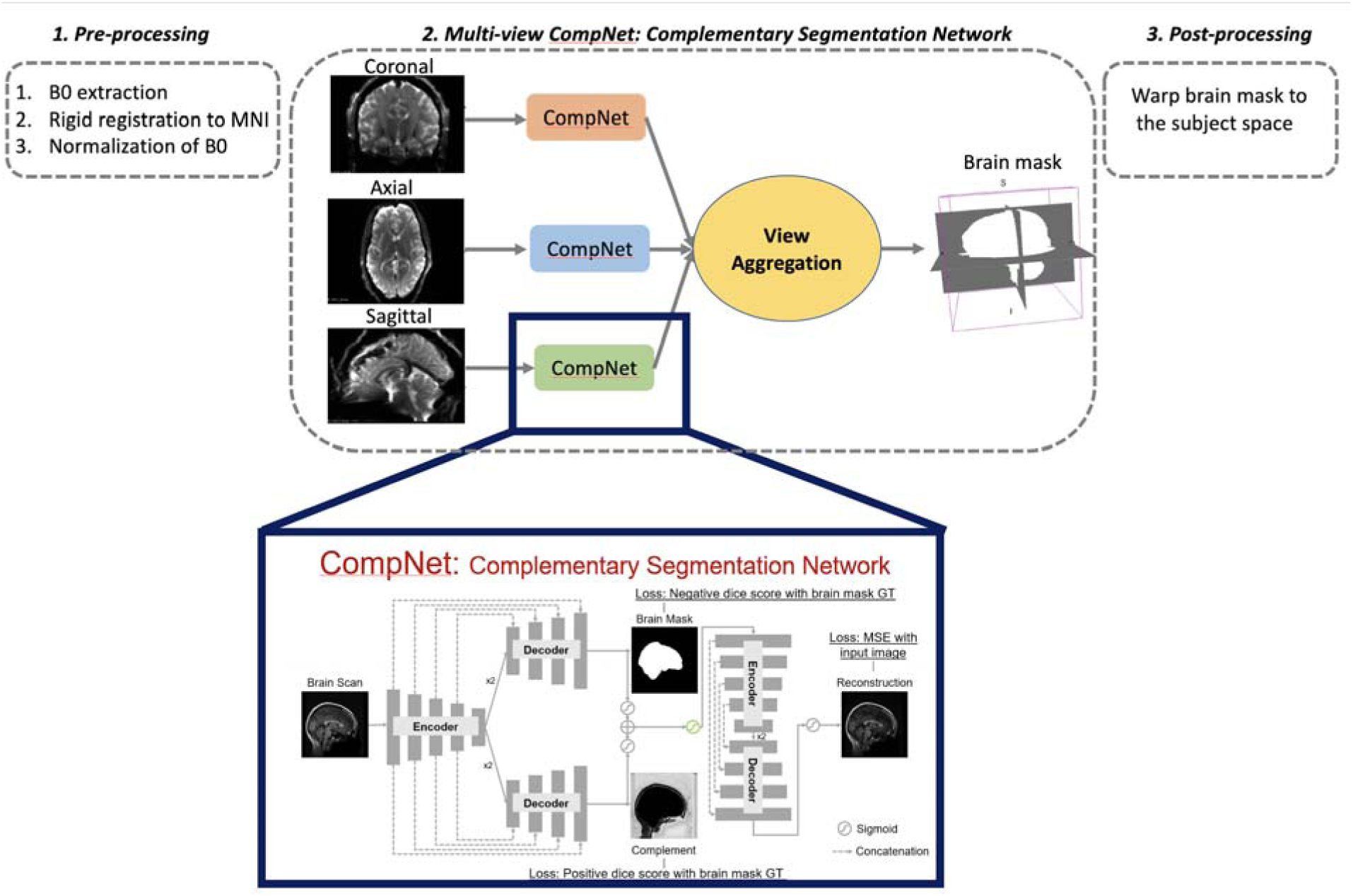
CNN DMRI Brain Segmentation: *(1) Pre-processing; (2) Multi-view CompNet; (3) Post-processing*. Multi-view CompNet includes three branches: (i) Segmentation Branch - learns the brain region; (ii) Complementary Branch - learns complement of the brain region; (iii) Reconstruction Branch - provides direct feedback to the segmentation and complementary branches and expects reasonable reconstructions. Segmentation and reconstruction branches include a series of encoder and decoder networks with a kernel of size 3 × 3. The number of convolutional filters in the encoder starts from 32, followed by 64, 128, 256, and 512, while the number in the decoder starts from 256, followed by 128, 64, and 32.

