## Supplementary Materials for "Harmonized diffusion MRI data and white matter measures from the Adolescent Brain Cognitive Development Study" (from another version of this preprint: version 1, 2023-04-04)

**Supplementary Material:**

| **Scanner** | **Site** | **Manufacturer** | **Manufacturer model** | **Device serial number** | **Software Version** |
| --- | --- | --- | --- | --- | --- |
| **1** | 1 | Philips Medical Systems | Achieva dStream | HASH6b4422a7 | 5.3.0\5.3.0.0 |
| **2** | 1 | Philips Medical Systems | Achieva dStream | HASH6b4422a7 | 5.3.1\5.3.1.0 |
| **3** | 1 | Philips Medical Systems | Achieva dStream | HASH6b4422a7 | 5.4.1\5.4.1.1 |
| **4** | 2 | SIEMENS | Prisma_fit | HASH1314a204 | syngo MR E11 |
| **5** | 3 | SIEMENS | Prisma | HASH5b0cf1bb | syngo MR E11 |
| **6** | 4 | GE MEDICAL SYSTEMS | DISCOVERY MR750 | HASH4b0b8b05 | 25\LX\MR  Software release:DV25.0_R02_1549.b |
| **7** | 4 | GE MEDICAL SYSTEMS | DISCOVERY MR750 | HASH4b0b8b05 | 27\LX\MR  Software release:DV26.0_R02_1810.b |
| **8** | 4 | GE MEDICAL SYSTEMS | DISCOVERY MR750 | HASHfeb7e81a | 25\LX\MR  Software release:DV25.0_R02_1549.b |
| **9** | 4 | GE MEDICAL SYSTEMS | DISCOVERY MR750 | HASHfeb7e81a | 27\LX\MR  Software release:DV26.0_R02_1810.b |
| **10** | 5 | SIEMENS | Prisma_fit | HASH311170b9 | syngo MR E11 |
| **11** | 6 | SIEMENS | Prisma_fit | HASH96a0c182 | syngo MR E11 |
| **12** | 7 | SIEMENS | Prisma_fit | HASH65b39280 | syngo MR E11 |
| **13** | 8 | GE MEDICAL SYSTEMS | DISCOVERY MR750 | HASH5b2fcf80 | 25\LX\MR  Software release:DV25.0_R02_1549.b |
| **14** | 8 | GE MEDICAL SYSTEMS | DISCOVERY MR750 | HASH5b2fcf80 | 27\LX\MR  Software release:DV26.0_R01_1725.a |
| **15** | 9 | SIEMENS | Prisma_fit | HASH4d1ed7b1 | syngo MR E11 |
| **16** | 10 | GE MEDICAL SYSTEMS | DISCOVERY MR750 | HASHd7cb4c6d | 25\LX\MR  Software release:DV25.0_R02_1549.b |
| **17** | 10 | GE MEDICAL SYSTEMS | DISCOVERY MR750 | HASHd7cb4c6d | 27\LX\MR  Software release:DV26.0_R01_1725.a |
| **18** | 10 | GE MEDICAL SYSTEMS | DISCOVERY MR750 | HASHe3ce02d3 | 27\LX\MR  Software release:DV26.0_R01_1725.a |
| **19** | 11 | SIEMENS | Prisma | HASH03db707f | syngo MR E11 |
| **20** | 12 | SIEMENS | Prisma_fit | HASH31ce566d | syngo MR E11 |
| **21** | 12 | SIEMENS | Prisma_fit | HASHe4f6957a | syngo MR E11 |
| **22** | 13 | GE MEDICAL SYSTEMS | DISCOVERY MR750 | HASH69f406fa | 27\LX\MR  Software release:DV25.1_R01_1617.b |
| **23** | 13 | GE MEDICAL SYSTEMS | DISCOVERY MR750 | HASH69f406fa | 27\LX\MR Software release:DV26.0_R02_1810.b |
| **24** | 13 | GE MEDICAL SYSTEMS | DISCOVERY MR750 | HASHc3bf3d9c | 25\LX\MR  Software release:DV25.0_R02_1549.b |
| **25** | 13 | GE MEDICAL SYSTEMS | DISCOVERY MR750 | HASHc3bf3d9c | 27\LX\MR  Software release:DV26.0_R02_1810.b |
| **26** | 14 | SIEMENS | Prisma | HASH11ad4ed5 | syngo MR E11 |
| **27** | 14 | SIEMENS | Prisma_fit | HASH7f91147d | syngo MR E11 |
| **28** | 15 | SIEMENS | Prisma_fit | HASH7911780b | syngo MR E11 |
| **29** | 16 | SIEMENS | Prisma | HASH3935c89e | syngo MR E11 |
| **30** | 17 | Philips Medical Systems | Achieva dStream | HASHdb2589d4 | 5.3.0\5.3.0.0 |
| **31** | 17 | Philips Medical Systems | Achieva dStream | HASHdb2589d4 | 5.3.0\5.3.0.3 |
| **32** | 17 | Philips Medical Systems | Achieva dStream | HASHdb2589d4 | 5.3.1\5.3.1.0 |
| **33** | 17 | Philips Medical Systems | Achieva dStream | HASHdb2589d4 | 5.4.0\5.4.0.1 |
| **34** | 17 | Philips Medical Systems | Achieva dStream | HASHdb2589d4 | 5.4.1\5.4.1.1 |
| **35** | 18 | GE MEDICAL SYSTEMS | DISCOVERY MR750 | HASHa3e45734 | 25\LX\MR  Software release:DV25.0_R02_1549.b |
| **36** | 18 | GE MEDICAL SYSTEMS | DISCOVERY MR750 | HASHa3e45734 | 27\LX\MR  Software release:DV26.0_R01_1725.a |
| **37** | 18 | GE MEDICAL SYSTEMS | DISCOVERY MR750 | HASHa3e45734 | 27\LX\MR  Software release:DV26.0_R03_1831.b |
| **38** | 19 | Philips Medical Systems | Ingenia | HASH5ac2b20b | 5.3.0\5.3.0.0 |
| **39** | 19 | Philips Medical Systems | Ingenia | HASH5ac2b20b | 5.3.1\5.3.1.0 |
| **40** | 19 | Philips Medical Systems | Ingenia | HASH5ac2b20b | 5.3.1\5.3.1.1 |
| **41** | 19 | Philips Medical Systems | Ingenia | HASH5ac2b20b | 5.3.1\5.3.1.2 |
| **42** | 20 | SIEMENS | Prisma | HASHd422be27 | syngo MR E11 |
| **43** | 20 | SIEMENS | Prisma_fit | HASHc9398971 | syngo MR E11 |
| **44** | 21 | SIEMENS | Prisma | HASH4036a433 | syngo MR E11 |
| **45** | 21 | SIEMENS | Prisma_fit | HASHb640a1b8 | syngo MR E11 |

**Supplementary Table 1.** 45 different scanners in 21 sites of the ABCD study.

| **Scanner** | **Site** | **Manufacturer** | **Manufacturer model** | **Device serial number** | **Software Version** |
| --- | --- | --- | --- | --- | --- |
| **4** | 2 | SIEMENS | Prisma_fit | HASH1314a204 | syngo MR E11 |
| **5** | 3 | SIEMENS | Prisma | HASH5b0cf1bb | syngo MR E11 |
| **6** | 4 | GE MEDICAL SYSTEMS | DISCOVERY MR750 | HASH4b0b8b05 | 25\LX\MR  Software release:DV25.0_R02_1549.b |
| **7** | 4 | GE MEDICAL SYSTEMS | DISCOVERY MR750 | HASH4b0b8b05 | 27\LX\MR  Software release:DV26.0_R02_1810.b |
| **8** | 4 | GE MEDICAL SYSTEMS | DISCOVERY MR750 | HASHfeb7e81a | 25\LX\MR  Software release:DV25.0_R02_1549.b |
| **9** | 4 | GE MEDICAL SYSTEMS | DISCOVERY MR750 | HASHfeb7e81a | 27\LX\MR  Software release:DV26.0_R02_1810.b |
| **10** | 5 | SIEMENS | Prisma_fit | HASH311170b9 | syngo MR E11 |
| **11** | 6 | SIEMENS | Prisma_fit | HASH96a0c182 | syngo MR E11 |
| **12** | 7 | SIEMENS | Prisma_fit | HASH65b39280 | syngo MR E11 |
| **13** | 8 | GE MEDICAL SYSTEMS | DISCOVERY MR750 | HASH5b2fcf80 | 25\LX\MR  Software release:DV25.0_R02_1549.b |
| **14** | 8 | GE MEDICAL SYSTEMS | DISCOVERY MR750 | HASH5b2fcf80 | 27\LX\MR  Software release:DV26.0_R01_1725.a |
| **15** | 9 | SIEMENS | Prisma_fit | HASH4d1ed7b1 | syngo MR E11 |
| **16** | 10 | GE MEDICAL SYSTEMS | DISCOVERY MR750 | HASHd7cb4c6d | 25\LX\MR  Software release:DV25.0_R02_1549.b |
| **17** | 10 | GE MEDICAL SYSTEMS | DISCOVERY MR750 | HASHd7cb4c6d | 27\LX\MR  Software release:DV26.0_R01_1725.a |
| **18** | 10 | GE MEDICAL SYSTEMS | DISCOVERY MR750 | HASHe3ce02d3 | 27\LX\MR  Software release:DV26.0_R01_1725.a |
| **19** | 11 | SIEMENS | Prisma | HASH03db707f | syngo MR E11 |
| **20** | 12 | SIEMENS | Prisma_fit | HASH31ce566d | syngo MR E11 |
| **21** | 12 | SIEMENS | Prisma_fit | HASHe4f6957a | syngo MR E11 |
| **22** | 13 | GE MEDICAL SYSTEMS | DISCOVERY MR750 | HASH69f406fa | 27\LX\MR  Software release:DV25.1_R01_1617.b |
| **23** | 13 | GE MEDICAL SYSTEMS | DISCOVERY MR750 | HASH69f406fa | 27\LX\MR Software release:DV26.0_R02_1810.b |
| **24** | 13 | GE MEDICAL SYSTEMS | DISCOVERY MR750 | HASHc3bf3d9c | 25\LX\MR  Software release:DV25.0_R02_1549.b |
| **25** | 13 | GE MEDICAL SYSTEMS | DISCOVERY MR750 | HASHc3bf3d9c | 27\LX\MR  Software release:DV26.0_R02_1810.b |
| **26** | 14 | SIEMENS | Prisma | HASH11ad4ed5 | syngo MR E11 |
| **27** | 14 | SIEMENS | Prisma_fit | HASH7f91147d | syngo MR E11 |
| **28** | 15 | SIEMENS | Prisma_fit | HASH7911780b | syngo MR E11 |
| **29** | 16 | SIEMENS | Prisma | HASH3935c89e | syngo MR E11 |
| **35** | 18 | GE MEDICAL SYSTEMS | DISCOVERY MR750 | HASHa3e45734 | 25\LX\MR  Software release:DV25.0_R02_1549.b |
| **36** | 18 | GE MEDICAL SYSTEMS | DISCOVERY MR750 | HASHa3e45734 | 27\LX\MR  Software release:DV26.0_R01_1725.a |
| **37** | 18 | GE MEDICAL SYSTEMS | DISCOVERY MR750 | HASHa3e45734 | 27\LX\MR  Software release:DV26.0_R03_1831.b |
| **42** | 20 | SIEMENS | Prisma | HASHd422be27 | syngo MR E11 |
| **43** | 20 | SIEMENS | Prisma_fit | HASHc9398971 | syngo MR E11 |
| **44** | 21 | SIEMENS | Prisma | HASH4036a433 | syngo MR E11 |
| **45** | 21 | SIEMENS | Prisma_fit | HASHb640a1b8 | syngo MR E11 |

**Supplementary Table 2.** After excluding the dMRI data of the Philips scanner sites, 18 sites with 33 scanners were present in the ABCD study.

**Exclusion criteria in the template creation process:**

We excluded the subjects matching the criteria defined below in the template creation process. We note that even though these subjects were excluded from the datasets that were used to learn the scanner-related measurement differences, they were included in the harmonization process.

*These subjects were excluded from all groups: the subjects who have/had cerebral palsy, tumor, stroke, TBI, aneurysm, hemorrhage, hematoma, epilepsy, seizure, gestational weight < 1200 g, birth complications, abnormal/bad quality MRI scans. Adolescents who were diagnosed with a substance disorder, as well as adolescents whose parents were diagnosed with schizophrenia, bipolar disorder, MDD, ASD, and ADHD, were excluded.*
